# Hippocampal cAMP-dependent synaptic potentiation, ERK-dependent immediate-early gene activation, and context-dependent fear conditioning in mice are linked through dependence on the guanine nucleotide exchange factor RapGEF2

**DOI:** 10.1101/2022.04.30.490162

**Authors:** Sunny Zhihong Jiang, Meishar Shahoha, Hugo A. Tejeda, Uri Ashery, Lee E. Eiden

## Abstract

The MAP kinase ERK is important for neuronal plasticity underlying associative learning, yet specific molecular pathways for ERK activation in hippocampus are still largely undetermined. RapGEF2 has emerged as a neuron-specific cAMP sensor that mediates ERK activation. We investigated whether RapGEF2 might also be required for cAMP-dependent ERK activation leading to synaptic potentiation, and how this involvement might be penetrant to hippocampus-dependent learned behavior. We demonstrate that conditional knockout of Rapgef2 in forebrain neurons, specifically in dentate gyrus and CA1 of the hippocampus, leads to an attenuation of context-dependent fear conditioning, but not of cue-dependent fear conditioning, in mice. RapGEF2 knockout is associated with a reduction in cAMP-dependent synaptic potentiation at two central hippocampal synapses-the entorhinal cortex-granule cell synapse and the CA3-CA1 synapse. Furthermore, cAMP-induced postsynaptic potentiation requires both RapGEF2 and activation of ERK. Induction of Egr-1/Zif268 (and pERK), but not of c-Fos, immediately following fear conditioning, was abolished in CA1 and detate gyrus, in the absence of RapGEF2 expression in these hippocampal regions, thus revealing a link between learning (conditioning) and molecular pathways activated during conditioned fear memory formation. Hence, we suggest that contextual fear conditioning is mediated via RapGEF2-dependent ERK activation and downstream induction of Egr-1, via an underlying mechanism of cAMP-dependent long-term potentiation at hippocampal synapses. Cyclic AMP-dependent GEFs have been genetically associated as risk factors for schizophrenia, a disorder associated with cognitive deficits. This study provides a functional link between one of these cAMP-dependent GEFs, RapGEF2, and cognitive processes involved in associative learning.

## INTRODUCTION

Cyclic adenosine monophosphate (cAMP) is a key second messenger mediating neuronal changes linked to altered behavior responses (Greengard, 2001; Kandel, 2001). Neurotransmitters such as catecholamines and serotonin cause cAMP elevation via ligand-activated guanine nucleotide exchange factors (GPCRs) coupled to Gs. Cyclic AMP acts as a *third* messenger in neurons after activation of adenylyl cyclases I and VIII by Ca2+ influx triggered by neuronal depolarization (Ferguson and Storm, 2004).

Several protein kinases important in cognition are activated after cAMP elevation (Schwartz, 1993; Shaywitz and Greenberg, 1999; Rosenblum et al., 2002). Establishing in molecular detail whether and how their activation depends upon cAMP has been difficult (Impey et al., 1999). Protein kinase A has been prominently implicated as a cognitive kinase activated by cAMP, with CREB as its downstream effector for at least some forms of synaptic plasticity (Kandel, 2001; Athos et al., 2002). The MAP kinase ERK also has a critical role in neuronal plasticity, however the mechanism of its activation by cAMP has remained unclear (Impey et al., 1998). The difficulty of disentangling ERK and CREB signaling, and their respective activation(s) by cAMP is illustrated by seminal reports on the relative role(s) of Rap (Morozov et al., 2003) and PKA (Eckel-Mahan et al., 2008) in mediating ERK-dependent aspects of long-term potentiation (LTP) important for hippocampal-dependent memory tasks. In fact at the intercellular level, several protein kinases (Frey et al., 1993; Sacktor et al., 1993; Lledo et al., 1995; Abel et al., 1997; Atkins et al., 1998; Impey et al., 1998) have been linked to various types of LTP (Collingridge and Bliss, 1995), thought to instatiate an important intercellular mechanism for encoding associative memory (Abraham et al., 2019). Forging molecular links between cAMP and ERK signaling, despite the apparent centrality of both for associative learning in rodents underlying spatial memory (maze guidance) and context-dependent responses (fear conditioning) (Adams and Sweatt, 2002) has been a recalcitrant problem in neurobiology.

Recently, two guanine nucleotide exchange proteins (GEFs), now called Epac1 and Epac2, were discovered that respond to cAMP by activation of the small GTPase Rap, potentially establishing a pathway for activation of ERK, via cAMP, independent of PKA (de Rooij et al., 1998; Kawasaki et al., 1998; Gloerich and Bos, 2010). However, subsequent analysis revealed that Epac is not a direct regulator of ERK, at least in neuroendocrine cells (Enserink et al., 2002). Subsequently, a neuron-specific GEF, RapGEF2, was identified as a cAMP effector that preferentially activates the MAP kinase ERK in PC12 and related neuroendocrine cells (Emery and Eiden, 2012; Emery et al., 2017).

A signaling matrix in which cAMP activates cognitive kinases through parallel PKA-dependent and PKA-independent pathways is an attractive construct for better understanding of the molecular events controlling neuronal plasticity. In fact, we have recently proferred evidence that dopamine signaling through the D1 GsPCR activates both CREB and ERK phosphorylation in D1 dopaminoceptive neurons (Jiang et al., 2017), and that eliminating RapGEF2 from D1 medium spiny neurons (MSN) of the nucleus accumbens curtails cocaine-induced locomotor sensitization and conditioned place preference (Jiang et al., 2021), the latter a type of learning previously shown to require ERK-dependent D1-MSN neuroplasticity (Girault et al., 2007).

In this report, we extend investigation of RapGEF2-dependent signaling to hippocampus-based neuronal plasticity and hippocampal-dependent associative learning. Full expression of cAMP-dependent long-term post-synaptic potentiation in both the Schaffer collateral and perforant pathways, induced by treatment of hippocampal slices with the adenylate cyclase agonist forskolin (Fsk), requires expression of RapGEF2. Deletion of RapGEF2 from the CA1 and dentate gyrus of the mouse results in specific impairment of context-dependent fear conditioning, and ERK activation after context-footshock pairing in a fear conditioning paradigm. The data argue for consideration of the role of RapGEF2 as a cAMP sensor required for hippocampal neuronal signaling that mediates the pairing of conditioned and unconditioned stimuli required for associative learning.

## MATERIALS AND METHODS

### Animals

Mice (wild-type or transgenic) on C57BL6J background were housed 2-5 per cage and acclimatized to 12-hour light/12-hour dark cycle with food and water *ad libitum*. Animal care was approved by the NIMH Institutional Animal Care and Use Committee (IACUC) and conducted in accordance with NIH guidelines. The floxed RapGEF2 mouse strain RapGEF2^fl/fl^ and RapGEF2 KO mouse strain Camk2a-cre^+/-^::RapGEF2^fl/fl^ (cKO) were generated as described previously (Jiang et al., 2017).

### Immunohistochemistry

Immunohistochemistry was conducted as previously described (Jiang et al., 2017) after animal perfusion with 4% paraformaldehyde. Briefly, mouse brains were sectioned by Vibratome at a 40 μm thickness. Free-floating sections were washed in TBS containing 0.5% Triton X-100 (TBST; 3 washes, 15 min), incubated at room temperature in blocking solution (10% normal goat or donkey serum in TBST; 1h), and then incubated in primary antibody diluted in blocking solution overnight at 4°C. The following day, sections were washed in TBST (3 washes, 15 min), incubated in the dark in Alexa 555-conjugated or Alexa 488-conjugated goat anti-rabbit-IgG (1:300; Millipore Sigma) for 2 h following primary antibody incubation. Sections were mounted in Vectashield (Vector Laboratories, Burlingame, CA). The primary antibodies used were rabbit anti-pERK (1:1500, Cell Signaling Technology, Danvers, MA), anti-c-Fos (1:5000, EnCor Biotechnology Inc., Gainesville, FL), anti-Egr-1 (15F7) (1:1000, Cell Signaling Technology, Danvers, MA) and rabbit anti-Rapgef2 (NNLE-2, custom-made by Anaspec, see ref. (Jiang et al., 2017)). Confocal images were obtained on a Zeiss LSM 510 confocal microscope at the National Institute of Neurological Disorders and Stroke Light Imaging Facility. Immunoreactive (IR) signals of RapGEF2, phospho-ERK, c-Fos and Egr-1 from different brain areas as indicated were quantified by NIH Image J using the mean gray values of integrated density after being converted to gray scale.

### Hippocampal slice preparation

The first step of acute hippocampal slice preparation was cardiac perfusion with ice-cold cutting solution which contained (in mM): 215 sucrose, 2.5 KCl, 20 glucose, 26 NaHCO_3_, 1.6 NaH_2_PO_4_, 0.5 CaCl_2_, 4 MgCl_2_, 4 MgSO_4_. Next, the hippocampi were isolated from the brain, and placed in an agar template to allow transverse sectioning at 400 μm thickness with a VT1000S vibratome (Leica). The resulting slices were immediately transferred to a solution composed of 50% Cutting solution and 50% recording solution (50-50 solution) for 30 minutes. The recording solution contained (in mM): 124 NaCl, 2.5 KCl, 26 NaHCO_3_, 1 NaH_2_PO_4_, 2 CaCl_2_, 1.3 MgSO_4_ and 10 glucose. All solutions were equilibrated with 95% O_2_ and 5% CO_2_. Next, the 50-50 solution was replaced by 100% recording solution and incubated for an additional hour at room temperature, after which the slices were considered as ready for experiments.

### Slice electrophysiology

Hippocampal slices were chosen for experiments after visualization using infrared differential interference contrast (IR/DIC) and ensured to be intact. Hippocampal slices were continuously perfused, in a submerged chanber, with fresh aCSF at a rate of 2 ml per min at 32°C. Electrical stimulations were induced by a monopolar electrode which delivered square-wave current pulses of 0.1 millisecond duration. The currents were generated by a stimulus isolator (Isoflex, AMPI) connected to a borosilicate glass pipette pulled by a pipette puller (Narishige) to a tip size of ~5 μm. fEPSPs were recorded using a patch pipette filled with 0.5M NaCl (2-3 MΩ) connected to an EPC10 amplifier (HEKA). For measurements in the hippocampal medial perforant pathway (MPP), the stimulating electrode was placed in the middle molecular layer of the DG near the vertex of the DG. The recording electrode was also placed at the middle molecular layer of the DG, but more laterally towards the tip of the upper blade. For measurements in the hippocampal Schaffer Collateral (SC) pathway, the stimulating was placed in the *stratum radiatum* of the CA1 subregion, near its border with the CA2 subregion. The recording electrode was also placed in the *stratum radiatum* of the CA1 subregion, but more downstream to the stimulating electrode, towards the subiculum. Stimulation intensity and electrode placement were calibrated to yield clean fEPSPs of an amplitude of 0.5-0.8 mV. fEPSPs were elicited by delivering paired pulses of 25 Hz once every 20 seconds. Before each experiment, a stable baseline of at least 20 minutes was ensured.

### Data acquisition and statistical analysis

Electrophysiology data was acquired using the PatchMaster software (Heka). The recorded files were then exported into CSV files using Igor Pro 6.22A (Wavemetrics), and further analyzed using custom-made MATLAB scripts. Specifically, the traces were first smoothed using the Savitzky-Golay method, in order to remove high-frequency noise. Next, unless stated otherwise, fEPSPs were calculated as the peak amplitudes of the responses and were normalized by transforming the values into percentages of the baseline response 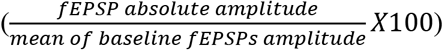. The facilitation was calculated using the paired-pulse ratio parameter which measures the ratio between the responses to two stimuli given in short order 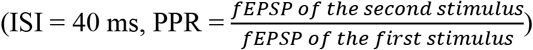.

Statistical analyses were done using Graphpad Prism 8 software. Results are reported as the mean ± SEM. Statistical significance was set to p<0.05 (marked by *; p<0.01 and p< 0.001 were signified using ** and ***, respectively). The statistical test used was repeated measures two-way ANOVA for most cases. All MATLAB scripts are available upon request.

### Reagents

Forskolin (Cat #1099), U0126 (Cat #1144) and bicuculline (Cat #0130) were purchased from Tocris Bioscience (Bristol, UK), and were used at concentrations of 25, 20 and 10 μM, respectively. U0126 incubation was begun 20 minutes before addition of forskolin and continued for 40 minutes. All working solutions contained ≤ 0.1% DMSO.

### Western blot

Mouse brain tissues from hippocampal CA1, CA3 and DG were dissected out in cold PBS and snap-frozen. Samples were sonicated on ice with RIPA buffer supplemented with Halt Protease and Phosphatase Inhibitor Cocktail (Thermo Scientific). RIPA-insoluble fractions were removed by centrifugation (3000 rpm for 10 min at 4°C). Supernatants were retained, and protein concentration was determined by MicroBCA Protein Assay kit (Thermo Scientific) according to the protocol provided by the manufacturer. Western blot was performed as described previously (Jiang et al., 2017). The rabbit polyclonal anti-GAPDH (D16H11) (Cell Signaling Technology, Cat# 5174; 1:1000) and Rabbit anti-RapGEF2 (NNLE-2, 1:1000) were used. Immunoreactive bands were visualized with a SuperSignal West Dura Chemiluminescent Substrate (Thermo Scientific), photographed with a ChemiDoc Imaging system, and quantified with NIH ImageJ.

### RNAscope in situ hybridization

In situ hybridization was conducted using the RNAScope Multiplex Fluorescent V2 Kit (Cat# 323100) (ACD Bio), as directed in the manual. Briefly, fresh-frozen mouse brains were sectioned coronally or sagittally on a cryostat to 12 μM. Sections were mounted on Superfrost Plus slides (Fisher Scientific). Slides were fixed with 4% paraformaldehyde for 20 minutes at 4^0^C, then dehydrated in an ethanol gradient. Hydrogen peroxide and protease treatments, probe hybridization and signal development proceeded as described in the manual. Probes were Mm-Fos (Cat No. 316921) and Mm-Egr1-C2 (cat No. 423371-C2). These were developed with Opal 520 and Opal 570 fluorophores (Akoya Biosciences). Confocal images were obtained on a Zeiss LSM 510 confocal microscope at the National Institute of Neurological Disorders and Stroke Light Imaging Facility.

#### Behavioral tests

Animal behavioral tests were conducted in NIMH and NICHD Rodent Behavioral Core Facilities.

### Home cage monitoring

Mice were monitored for locomotor activity for two days in individual home cages under normal vivarium conditions using the Photobeam Activity System-Home Cage (PAS-HC) (San Diego Instruments, San Diego, CA, USA).

#### Elevated zero maze

The elevated zero maze for anxiety was performed by placing the mouse into an open quadrant of a continuous circular 5.5 cm-wide track elevated 65 cm above the floor and divided into alternating walled and open quadrants in a dimly lit room, for 6 min. Video recording from above in the last 5 min was automatically scored for time spent in each quadrant (Top Scan software suite, Clever System Inc.).

#### Open field

Mice were placed in a Photobeam Activity System-open field (PAS-OF) (20” X 20” frame dimension, 16 X 16 photo beam configuration) to monitor locomotor activity for 1 hour in a novel environment.

#### Rotarod

Rotarod test was performed using an Accurotor rotarod (AccuScan Instruments). Animals were tested concurrently in four separate 11 cm-wide compartments on a rod approximately 3 cm in diameter and elevated 35 cm. The apparatus started at an initial speed of 4 rpm and gradually accelerated at a rate of 0.2 rpm/s. The latency to fall from the rod was recorded with a cut-off time of 2 min. Each animal was assessed over ten trials with 20-min intertrial intervals.

#### Hot Plate

Pain sensitivity of mice was tested using a Hot Plate Analgesia Meter (San Diego Instruments, San Diego, CA, USA). Mice were placed on the hot plate (55±0.1^0^C). The mice were constrained to the hot plate by clear acrylic walls (19 cm tall, open top). The latency to respond with either hind paw lick, hind paw flick, or jump was measured, and the mouse was immediately removed from the hot plate and returned to its home cage. The cut-off time was 30 s. Animals were tested individually and were not habituated to the apparatus prior to testing.

#### Barnes Maze

A modified Barnes maze (Patterned Barnes maze, PBM) (Barnes, 1979; Faizi et al., 2012) was used to test spatial learning and memory. The PBM consists of a 122 cm diameter circular platform with a height of 91.44 cm. The platform consists of 40 holes; 16 holes in the outer ring, 16 holes in the middle ring and 8 holes in the inner ring. The holes are of diameter 10 cm. A plastic escape tube with a ramp was positioned beneath one of the holes to serve as the escape box. Distal extra-maze visual cues were placed on the walls of the room. The high overhead lighting at 1200 lux was used to create aversive motivation for the subjects. Trials were video recorded and scored by ANY-maze software (ANY-maze, Wood Dale, IL, USA) for measures including latency to find the hidden platform, total distance traveled, and speed. The animals were habituated in the escape box for 60 sec one day before training. Training trials included 7 days of training (two trials for each animal daily with 30 min of intertrial interval and 3 min cutoff time for each trial). Animals took a rest for two days and had a probe test on day 10. In the probe trial, the escape box was removed from the target hole and animals were allowed to explore on the platform for 3 mins.

#### Fear conditioning

On day 1 of the experiment, mice were habituated to the test chamber (Med Associates) for 5 min in context A which was used for conditioning, then 5 min in context B which is distinct from context A. Context A consisted of a square metal chamber, shock grid floor and external chamber light on one side. Context A was cleaned between each trial using 70% ethanol and the testing room was dimly lit. Context B consisted of patterned paper walls, plastic chamber floor and an internal blue light on top of the chamber. Context B was cleaned with household glass cleaner between trials. The testing room was dark for Context B. On day 2, the mouse was exposed to context A for 2.5 min, followed by 1 pair of a tone (3000 Hz, 85 dB, 30 s) with a foot shock (0.5 mA, 2s). Contextual fear memory was tested at 3 and 24 h after fear conditioning by placing the mouse in the training chamber for 3 min. Cued fear memory was tested at 24 h after fear conditioning by exposing the mouse to a tone (3000 Hz, 85 dB, 30s) following free running for 2.5 min in context B. Freezing was identified by complete immobility (at least 2s) with the Video Freeze 4 software (Med Associates).

### Experimental design and statistical analysis

Mice of both sexes were used in all studies. The sample size (n) per group is indicated in the figure legends for each experiment. Statistical analyses were conducted using Sigma Plot 14.0 (Systat Software).

Student’s t-tests and factorial model analysis of variances (ANOVA) were employed where appropriate. *Post hoc* analyses were performed using the Bonferroni Test. Data were reported using histograms and scatter plots to represent mean ± s.e.m and individual data in each group. Differences were considered to be significant when p<0.05.

## RESULTS

### Ablation of RapGEF2 protein expression in hippocampal CA1 and DG

To identify the role of RapGEF2-dependent signaling in associative learning, we generated a mouse line that permitted us to selectively knockdown RapGEF2 in forebrain projection neurons. To this end, we used the Cre/loxP system, by crossing Camk2α-Cre and floxed RapGEF2 mouse lines (Jiang et al., 2017). In these mice, exon 4 of the mouse RapGEF2 gene, and RapGEF2 protein, were eliminated in corticolimbic areas of mouse forebrain, including medial prefrontal cortex (mPFC), and basolateral amygdala (BLA), as previously reported (Jiang et al., 2021). In the current study we extended our investigations to the hippocampus and revealed that RapGEF2 protein expression was ablated in mouse hippocampus in an age-dependent manner. RapGEF2 immunostaining of brain sections from Camk2a-Cre^+/-^::RapGEF2^fl/fl^ (cKO) mice showed a ~50% reduction in RapGEF2 protein level in hippocampal CA1, but not in DG and CA3, at the age of postnatal 5 weeks (Figure 1A, left panel). RapGEF2 protein was largely ablated in both CA1 and DG areas by age 10 weeks (Figure 1A, right panel; 1B, Student’s t-test, *p*<0.001), with 11.83 ± 4.28% remaining in CA1 and 11.43 ± 2.68% remaining in DG of cKO hippocampus based on quantification of immunohistochemical staining for RapGEF2. No reduction was observed in the CA3 region. Similar results were obtained by western blot using protein lysates from subregions of the hippocampus (Figure 1A, right panel; 1B, Student’s t-test, *p*<0.001). Only 18.13 ± 3.75% of RapGEF2 protein remained in CA1 and 21.03 ± 5.63% remained in DG in cKO hippocampus (Figure 1C). Accordingly, we used mice older than 10 weeks for all further experiments.

**Figure 1.**
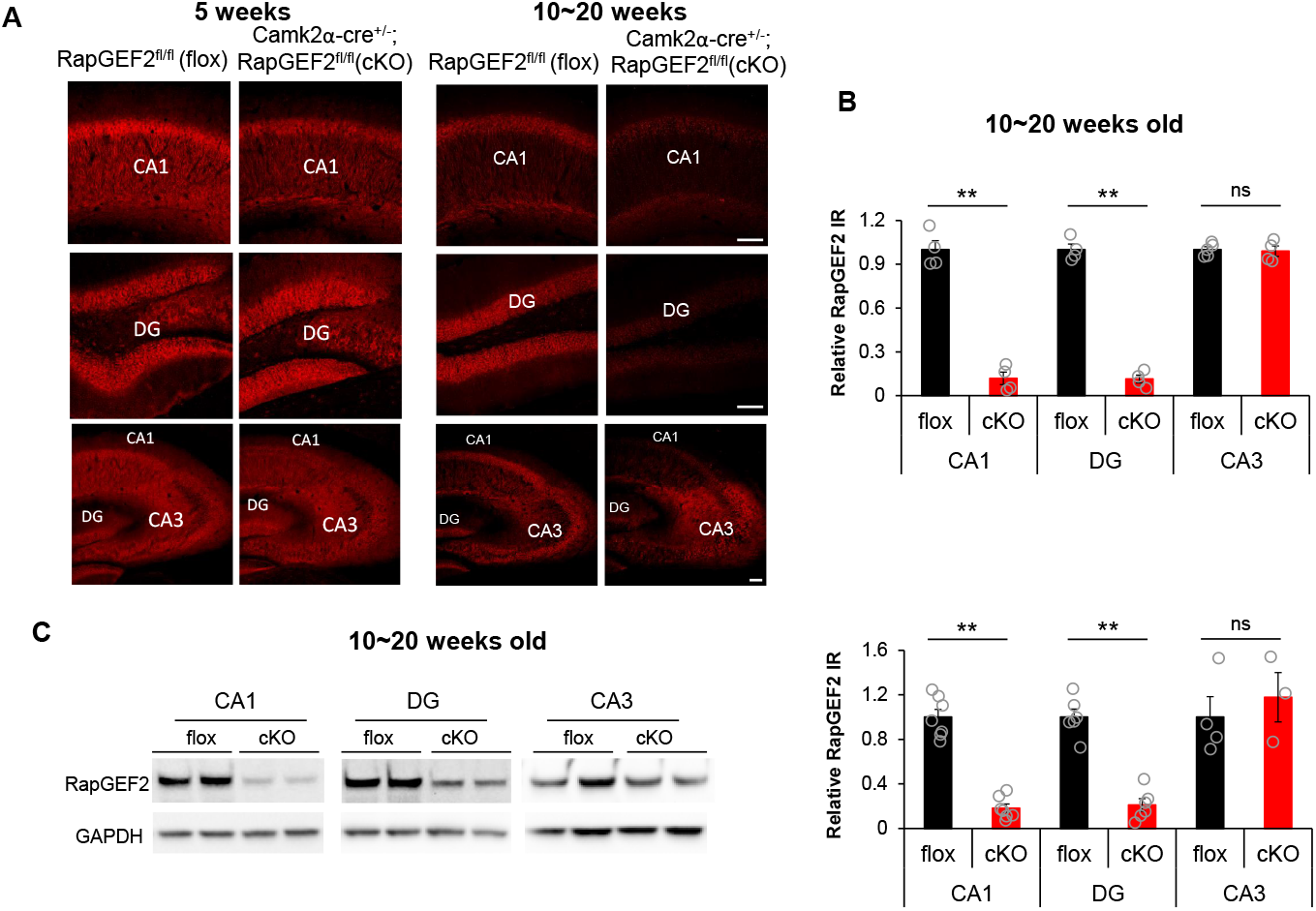
Ablation of RapGEF2 protein expression in hippocampal CA1 and DG. (A) Representative immunohistochemical images with RapGEF2 antibody (NNLE-2) for hippocampal subregions (CA1, DG and CA3) from flox and cKO mice at 5-weeks old (left panel) and 10~20 weeks old (right panel). Scale bar: 100μm. (B) RapGEF2 immunoreactive (IR) signals in hippocampal CA1, DG and CA1 from cKO mice (10~20 weeks old) were quantified with NIH Image J and compared to RapGEF2 IR signals from flox mice. The result indicated a significant reduction in RapGEF2 levels in the CA1 and DG hippocampal subregions, but not in the CA3 subregion. N=4~5 for each group. Student’s t-test for each region, \*\**p*<0.001. (C) Western blots using protein lysates from hippocampal subregions of flox and cKO mice showed similar results that RapGEF2 was downregulated in CA1 and DG. N=3~7 for each group, Student’s t-test for each region, \*\**p*<0.001.

### Impairment of contextual fear memory consolidation following RapGEF2 protein ablation in hippocampus

We first assessed whether RapGEF2 ablation might induce deficits in primary behavioral responses unrelated to learning and memory, but potentially influencing them. Accordingly, mice were subjected to serial behavioral tests that examine locomotor activity, innate anxiety-like behavior, and motor coordination. Long-term locomotor activity (48 h) was monitored in cKO mice and their floxed controls in home cages. The two genotypes showed similar activity patterns during day and night (Figure 2A_1_): animals were more active at night, indicated by the overall increased mobile behaviors at night compared with that during daytime. No difference was observed between floxed and cKO mice in the total number of beam breaks during 48 h (Figure 2A_1_, Student’s t-test, T=0.252, P=0.803). Exploratory behavior in a novel environment was assessed in an elevated zero maze (Figure 2A_2_) and an open field (Figure 2A_3_). Both genotypes preferred the closed arm when they were allowed to explore on the elevated zero maze for 5 min. As shown in Figure 2A_2_, cKO mice spent 22.22±1.28% of their time in open arms, similar to their floxed controls (19.77±1.21%)(Student’s t-test, T=1.389, P=0.172). This suggested that there was no change in anxiety-related behavior for animals with RapGEF2 ablation in the hippocampus. cKO mice and floxed control mice also showed similar exploratory behavior and habituation curves in a novel open field (Figure 2A_3_). A rotarod test was performed to examine the motor coordination of the animals. No significant differences between floxed and cKO mice in learning curves during the rotarod were observed; characterized by the increased latency to fall, across 10 trials (Figure 2A_4_, Two-way Repeated Measures ANOVA, training trial effect F_(9, 405)_=26.24, p<0.001; genotype effect F_(1,405)_=0.401, p=0.53; genotype X trial interaction F_(9, 405)_=0.244, p=0.988; trial 10 vs trial 1, p<0.001 for both floxed and cKO mice). To further determine whether RapGEF2 cKO selectively impaired hippocampus-dependent associative learning, we tested floxed controls and cKO mice in the Patterned Barnes Maze (PBM) to examine hippocampus-dependent spatial learning and memory. In this test, the animal was placed on a platform and learned to find an escape box (Figure 2B_1_) using visual cues outside the maze. Over seven days of training, learning curves for floxed controls and cKO mice were similar, with no difference in escape latency between groups (Figure 2B_2_, Two-way Repeated Measures ANOVA, genotype effect: F_(1, 222)_=0.00667, p=0.935; training day effect F_(6, 222)_=29.634, p<0.001; genotype X day interaction F_(6, 222)_=0.455, p=0.841. Day 7 vs Day 1, p<0.001 for both floxed and cKO mice). Animals’ behaviors in the probe trial also supported normal spatial learning for both groups (Figures 2B_3_ and 2B_4_): Floxed control and cKO mice showed preferential searching in zone 4, where the escape box was located during training trials. Two-way ANOVA followed by *post hoc* Bonferroni t-test showed that there was a significant zone effect (F_(3,148)_=288.714, p<0.001; zone 4 vs other zones, p<0.001), but no genotype effect (F_(1,148)_=5.21E-0.6, p=0.998). Escape latencies in the probe trial did not differ between floxed and cKO mice (Student’s t-test, T=277, P=0.077). Results above indicated that RapGEF2 cKO mice show normal locomotor activity, exploration, balance and motor coordination, and spatial navigation and memory.

**Figure 2.**
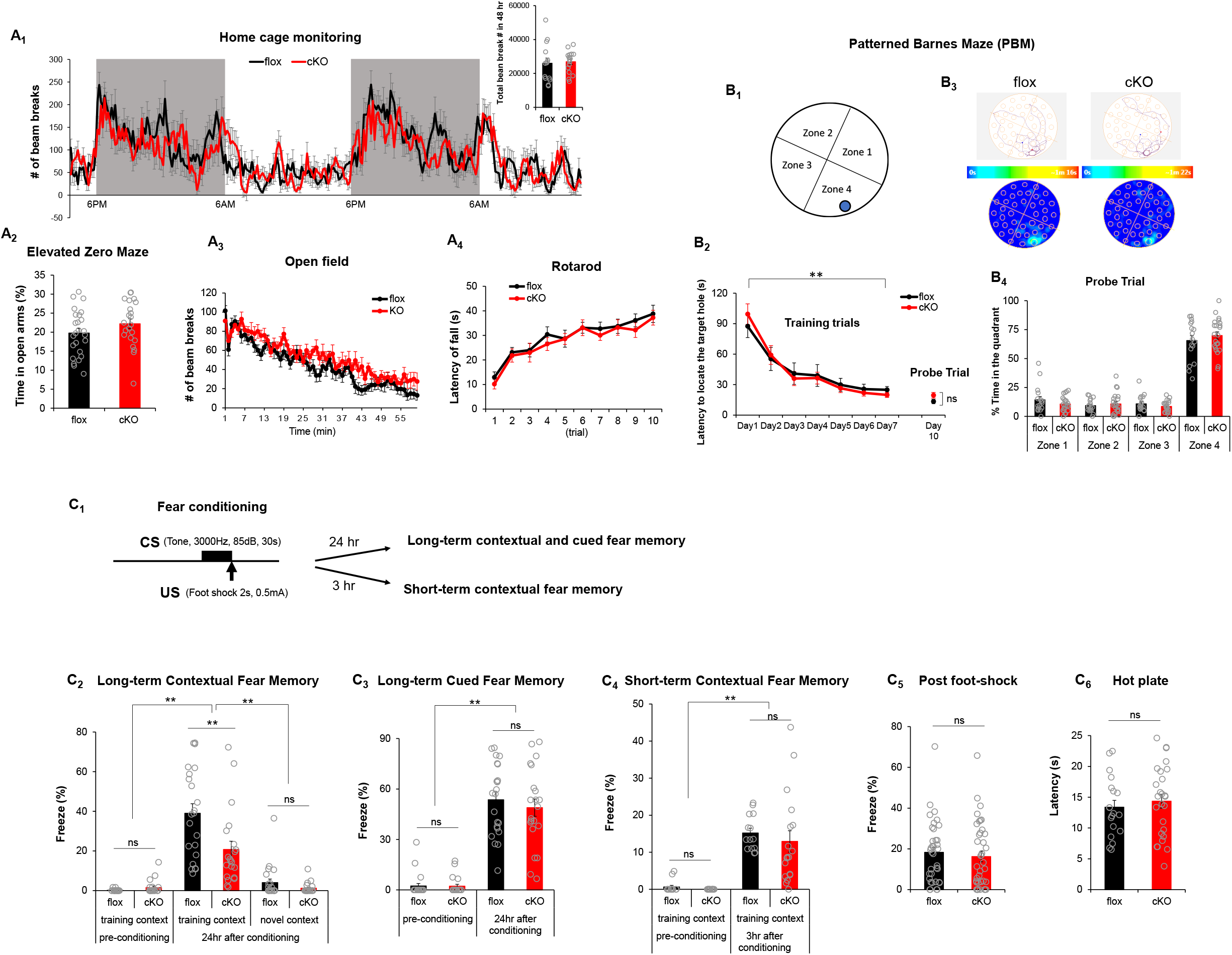
Camk2a-cre^+/-^::RapGEF2^fl/fl^ (cKO) mice show deficit in consolidation of contextual fear memory, with normal locomotor activity, exploration, balance and motor coordination, pain sensitivity and spatial navigation and memory. (A_1_~A_4_) Camk2a-cre^+/-^::RapGEF2^fl/fl^ (cKO) mice show normal locomotor activity, exploration, balance and motor coordination. The Long-term locomotor activity for flox and cKO was monitored in their home-cage for 48 hr. The locomotive behaviors of flox and cKO are not different, showing similar pattern during day and night in their familiar home-cage (A_1_, N=14~16 for each group). cKO mice were not more anxious than control animals in the elevated zero maze, as shown by the similar staying time in unprotected open arms (A2, N=22~25 for each group). The exploration behaviors were tested in a novel open field. cKO mice showed similar locomotor activity and habituation curve compare to flox mice (A3, N=10 for each group). A rotarod test was performed to examine the motor coordination. The results demonstrated no significant difference between flox and cKO mice (A_4_, N=22~25 for each group). (B_1_~B_4_) Spatial learning and memory of cKO mice were not impaired in a Patterned Barnes Maze (PBM) test. The plateform of the maze had 40 holes and were divided into 4 zones (B_1_). cKO and flox controls showed similar learning curve, indicated by similar latency to locate the target hole on each day (B_2_). Each animal ran two trials daily, and the average time to find the escape box were plotted for the latency on each day. In the probe trial, both flox and cKO mice spent more time to explore zone 4 (with target hole) indicated by the navigation map and the staying heat map (B_3_). cKO mice spend similar time to locate the target hole, as flox controls (B4). N=17 ~22 for each group, two-way repeated measures ANOVA, **p<0.001, day 1 vs day 7. (C_1_~C_6_) Consolidation of contextual fear memory was impaired in Camk2a-cre^+/-^::RapGEF2^fl/fl^ mice. The scheme of a fear conditioning test used for cKO and flox control mice (C_1_). cKO mice showed impaired contextual memory to the training context, not novel context, 24 h later compared to controls (C_2_). However, both flox and cKO mice showed normal levels of freezing during the tone presentation in a non-training context when memory was retrieved 24 hr after conditioning (C_3_, two-way ANOVA following by *post hoc* Bonferroni t-test, **p<0.001, N= 22 for floxed mice, N=21 for cKO mice). Contextual freezing 3 hr after the conditioning was similar between cKO and flox mice, suggesting acquisition and retrieval of memory was not affected in cKO mice (C_4_, two-way ANONA following by *post hoc* Bonferroni t-test, **p<0.001, N=14 for flox group, N=18 for cKO group). Flox and cKO mice showed similar freezing level immediately after foot-shock during fear conditioning (C_5_, N=36 for floxed, 39 for cKO) and similar latency in a hot plate test (C_6_, N=19~26 for each group), suggesting no differences in pain sensitivity between the two groups of mice.

It is well established that contextual fear conditioning is hippocampus-dependent, while both contextual and cued fear conditioning require the BLA (Phillips and LeDoux, 1992). To determine whether RapGEF2 cKO mice had deficits in contextual and/or cued fear conditioning, animals underwent fear conditioning procedures with both contextual and auditory components, wherein animals learned to associate a tone (85dB, 3KHz, 30 sec, co-terminated with foot shock) with an electric foot shock (0.5mA, 2s) in a chamber within a specific training context (Figure 2C_1_). Contextual fear memory was tested 24 h after fear conditioning by exposing mice to the training chamber or a novel chamber. Two-way ANOVA followed by *post hoc* Bonferroni t-test (Figure 2C_2_) indicated that there was a significant difference between floxed control and cKO mice when the contextual memory was retrieved 24 hours later (genotype effect: F_(1, 82)_=6.828, p=0.011; floxed vs cKO, p<0.001 in the training context 24 hr later). Both floxed and cKO mice could distinguish between training and novel contexts (Figure 2C_2_, Two-way ANOVA followed by *post hoc* Bonferroni t-test, context effect: F_(1, 82)_=66.802, p<0.001). Although RapGEF2 is ablated in BLA in cKO mice (Jiang et al, 2017, 2021), these mutant mice associate tone with the foot shock as did the floxed control mice, and did not differ from floxed controls in cued fear memory when tested 24 h after fear conditioning (Figure 2C_3_, Two-way ANOVA followed by *post hoc* Bonferroni t-test, to compare floxed and cKO mice during tone presentation. Conditioning effect: F_(1, 82)_=189.502, p<0.001; genotype effect: F_(1, 82)_=0.462, p=0.499; flox vs cKO after conditioning, p=0.351). Attenuation of contextual freezing 24 hr after fear conditioning in cKO mice was not due to impaired acquisition or retrieval of memory, because contextual freezing 3 hr after the conditioning was similar between cKO and floxed control mice (Figure 2C_4_, two-way ANOVA following by *post hoc* Bonferroni t-test, conditioning effect F_(1,60)_=62.085, p<0.001; genotype effect F_(1,60)_=0.703, p=0.405; flox vs cKO 3hr after condition, p=0.358). Flox and cKO mice showed similar freezing level immediately after foot-shock during fear conditioning (Figure 2C_5_, Student’s t-test, t=0.580, P=0.564) and similar latency in a hot plate test (Figure 2C_6_, Student’s t-test, T=-0.601, P=0.551), suggesting no differences in pain sensitivity between two groups. In summary, the fear conditioning results above indicated that hippocampus-dependent contextual fear memory consolidation was selectively impaired in Camk2a-Cre^+/-^::RapGEF2^fl/fl^ mice.

### ERK activation in hippocampus during fear conditioning is RapGEF2-dependent

We have recently identified RapGEF2 as a cAMP sensor/effector that links Gs-coupled GPCRs to the MAP kinase ERK in neuronal and endocrine cells in adult mammals (Emery et al., 2013; Emery et al., 2014; Jiang et al., 2017; Zhang et al., 2021). It has been previously demonstrated that pERK induction in hippocampus occurs during shock associated with context, but not in animals exposed only to the context, nor to animals shocked immediately without contextual paired experience (Atkins et al., 1998; Sindreu et al., 2007). To investigate whether RapGEF2 contributed to contextual fear memory consolidation via ERK signaling, hippocampal CA1 and DG sections were collected from floxed control and cKO mice at different time points (10 min, 30 min, or 60 min) after fear conditioning, and immunostained with phospho-ERK antibody (Figure 3). Sections from animals kept in the home cage (without fear conditioning, NFC) were used as controls. Two-way ANOVA followed by *post hoc* Bonferroni t-test was assessed to compare flox vs cKO at different time points. ERK activation in hippocampal CA1 region (Figure 3A & 3C) showed significant genotype effect (F_(1, 25)_=11.448, p=0.002), time effect (F_(3, 25)_=33.315, p<0.001) and genotype X time interaction (F_(3, 25)_=4.509, p=0.012). Floxed mice showed a significant increase in pERK in CA1 at 10 min or 30 min following fear conditioning (p<0.001, FC 10 min or FC 30min vs NFC), however, pERK levels at 60 min dropped to similar levels as home cage controls (p=1, FC 60 min vs NFC). cKO mice showed attenuation of pERK activation 10 min (p<0.001, cKO vs flox) or 30 min (p=0.011, cKO vs flox) after fear conditioning. Similarly, ERK activation in hippocampal DG region (Figure 3B & 3D) also showed a significant genotype effect (F_(1, 28)_=13.802, p<0.001), time effect (F_(3, 28)_=4.238, p=0.0014) and genotype X time interaction (F_(3, 28)_=3.572, p=0.026). There was an increase of pERK in DG at 10 min (p=0.008, FC 10 min vs NFC) or 30 min (p=0.016, FC 30 min vs NFC) following fear conditioning for floxed control mice, while cKO mice showed attenuation of pERK activation at these time points (cKO vs flox at FC 10 min, p<0.001; at FC 30 min, p=0.029).

**Figure 3.**
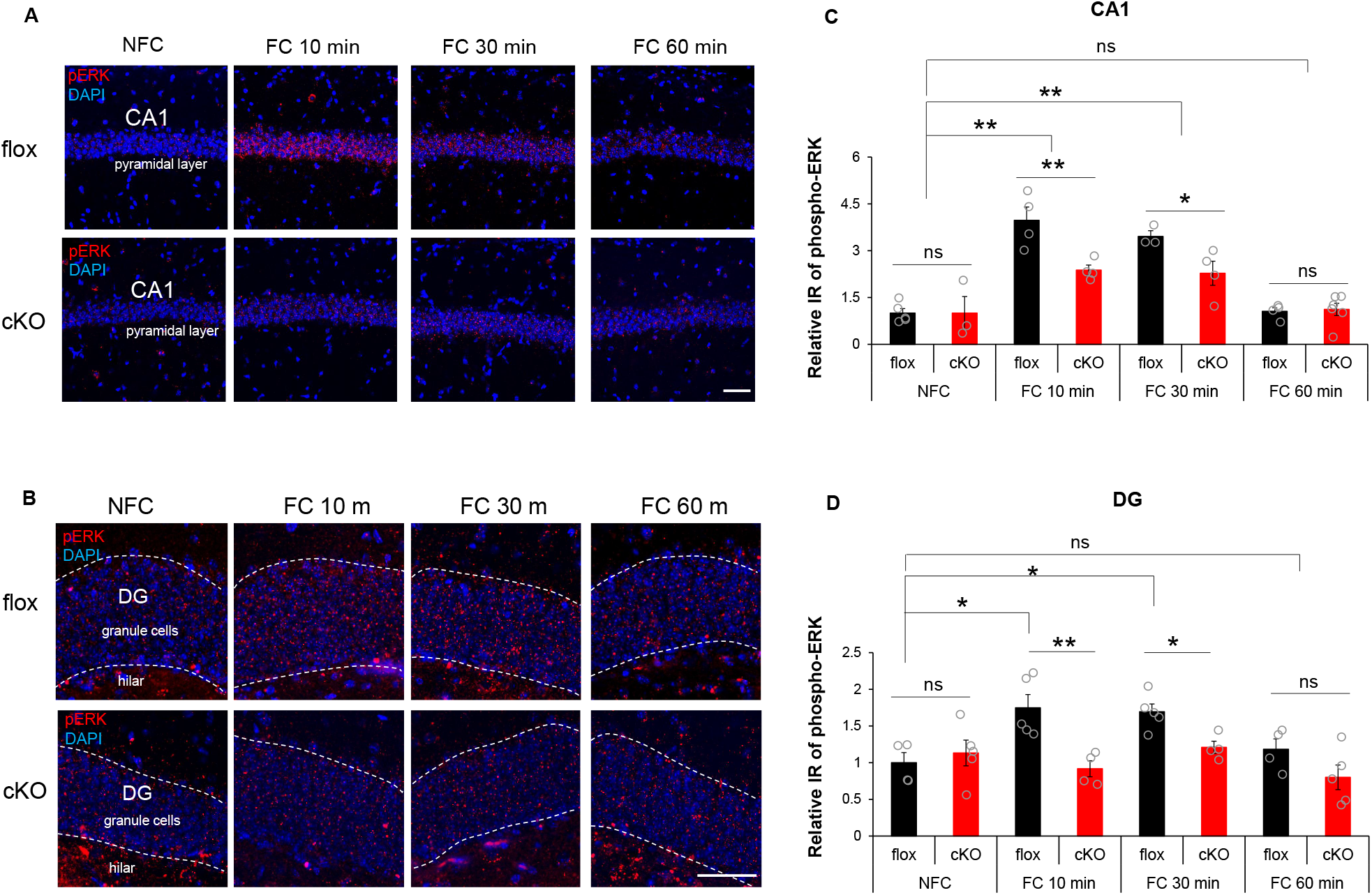
ERK activation in hippocampal subregions CA1 and DG following fear-conditioning. (A) & (B) Representative images of phospho-ERK staining (in red) in hippocampal CA1 pyramidal cell layer (A) or hippocampal DG granule cell layer (B) of flox and cKO mice 10 min (FC10 min) or 30 min (FC 30 min) or 60 min (FC 60 min) after fear conditioning or without fear conditioning (NFC). Scale bar: 50 μm. (C) & (D) Quantification of phospho-pERK IR in the CA1 (C) or DG (D) of flox and cKO mice at different time points after fear conditioning, showed RapGEF2-dependent activation of ERK 10 and 30 min after training. N=3~5 for each group. Two-way ANOVA following by *post hoc* Bonferroni t-test, *p<0.05; **p<0.001.

### Basal synaptic properties and short-term plasticity are intact in CaMKII-Rapgef2-cKO mice in the Schaffer collateral and the medial perforant pathways

RapGEF2-dependent signaling may modify the function of excitatory synapses within hippocampal circuitry that is essential for contextual fear conditioning. To determine if basal synaptic properties of hippocampal neurons were modified by the absence of RapGEF2 expression, we utilized ex vivo slice electrophysiology. We focused on two primary hippocampal synapses; the CA3-CA1 synapse through the Schaffer collateral (SC) pathway, and the entorhinal-cortex to DG synapse through the medial perforant pathway (MPP). Each of these hippocampal synapses exhibits robust synaptic plasticity. and has been implicated in generation of fear-related memories in contextual fear conditioning procedures. We first generated input-output (I/O) curves measuring the relationship between the electrical stimulation intensity and the amplitude of the resulting electrically-evoked fEPSP in the SC and MPP synapses. No differences between the I/O curves of cKO and floxed control slices were observed, either in SC (Figure 4A_1_, two-way ANOVA, F_(1,128)_ = 2.337, p = 0.1288) or MPP synapses (Figure 4A_2_, two-way ANOVA, F_(1,80)_ = 2.067, p = 0.1544). This suggests that SC and MPP synapses have similar baseline synaptic efficacies in cKO mice relative to their floxed controls. Changes in STP are indicative of presynaptic properties such as neurotransmitter release probability (P_r_). Thus, we tested if Rapgef2 depletion alters presynaptic release probability and short-term plasticity (STP) in the SC and MPP synapses. For this purpose, we performed measurements of paired-pulse ratios (PPR) obtained at different inter-spike intervals (ISIs used; 10, 20, 50, 100 and 200 milliseconds). Our results show that PPR values measured from the SC and MPP synapses did not show any genotype-dependent differences in PPR values, suggesting similar basal Pr of cKO and floxed controls (Figure 4B_1_, two-way ANOVA, F_(1,80)_ = 0.9947, p = 0.3216 and Figure 4B_2_, two-way ANOVA, F_(1,80)_ = 0.2109, p = 0.6473). In addition, we tested the facilitation pattern associated with a burst of four stimuli given with an ISI of 40 milliseconds. Significant facilitation or depression are seen following such stimulation in the SC and MPP, respectively. Consistent with the aforementioned results, cKO and floxed controls showed a similar burst facilitation/depression pattern (Figure 4C_1_ two-way repeated-measure ANOVA, F_(1, 15)_ = 0.06109, p = 0.8081 and Figure 4C_2_, two-way repeated-measure ANOVA, F_(1, 16)_ = 0.2387, p = 0.6317). Lastly, we compared all PPR values measured during the baseline phase of the above experiments and compared them between slices obtained from cKO and floxed littermates. We found no significant differences in basal PPR values between these two groups (Figure 4D_1_, Unpaired student’s t-test, p = 0.8479 and Figure 4D_2_, Unpaired student’s t-test, p = 0.8483). Overall, these results suggest that cKO and floxed control mice are identical in their basal synaptic and short-term plasticity properties.

**Figure 4.**
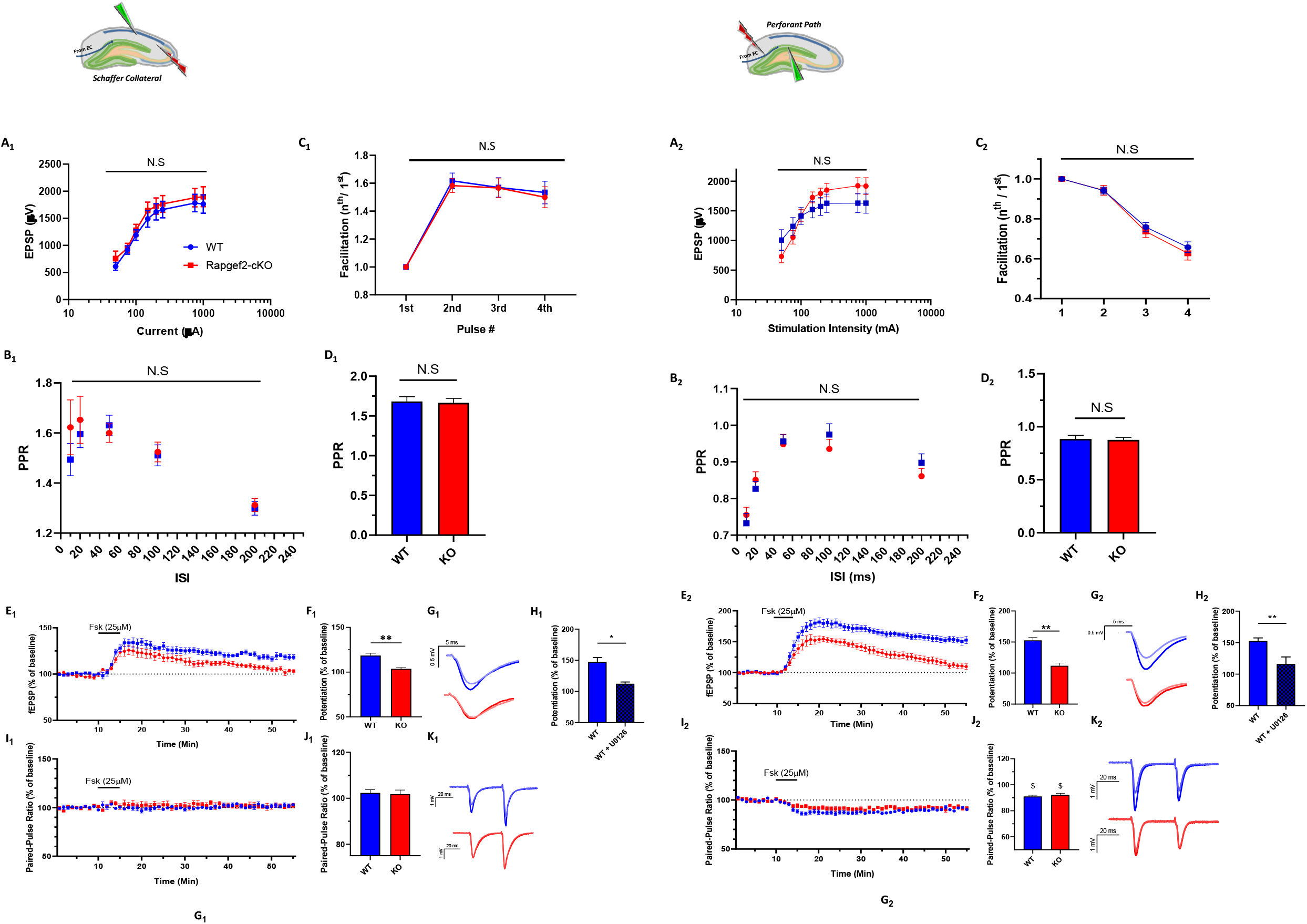
CaMKII-Rapgef2-cKO mice show impaired Fsk-induced potentiation without changes in STP in the SC-CA1 and MPP pathways. Results are shown symmetrically for the SC (1) and MPP (2) pathways A_1,2_) Input/Output (I/O) curves obtained from WT (floxed) (blue trace) and CaMKII-Rapggef2-cKO (red trace) mice show no differences in I/O curves (N=8 slices, 4 mice per group. Two-way ANOVA, p > 0.05 B_1,2_). Results from a comparison of PPR values at different inter-spike intervals (ISIs) suggest similar PPR between Rapggef2-cKO and WT (floxed) in both SC and MPP pathways (B_1_ and B_2_, respectively; N=7-11 slices, 4-6 mice, per group. Two-way repeated-measures ANOVA, p > 0.05 C_1,2_) Facilitation/depression pattern that results from a burst of 4-stimulations given at 25 Hz is not significantly different between Rapggef2-cKO and WT (floxed) in both SC and MPP pathways. (C_1_ and C_2_, respectively; N=7-11 slices, 4-6 mice, per group. Two-way ANOVA, p > 0.05 D_1,2_). No differences also in the baseline PPR values obtained during subsequent experiments. (D_1_ and D_2_, ISI = 40 milliseconds. N=9-14 slices, 5-7 mice, per group. Two-way ANOVA, p > 0.05). E_1,2_) fEPSP measurements before and after Fsk application. WT (floxed) littermates (blue) display bigger and long lasting Fsk-induced potentiation for SC and MPP synapses (E_1_ and E_2_, respectively) ccompare to Rapgef2-cKO (red). (N=4-10 slices, 3-5 mice per group. Two-way repeated-measures ANOVA, p = 0.0296 (SC) and p = 0.0001 (MPP). F_1,2_) Summary results of fEPSPs show a significant impairment of Fsk-induced potentiation in slices obtained from CaMKII-Rapgef2-cKO animals, unpaired student’s t-test, p = 0.1236 (SC), p < 0.0001 (MPP) G_1,2_). Representative traces of fEPSPs before (faint color) and 40 minutes after (bold color) Fsk administration for WT (floxed; blue) and CaMKII-Rapgef2-cKO (red) conditions. H_1,2_). Results from comparing WT (floxed) slices that were applied with Fsk only, and slices that had a prior preincubation with U0126 (20μM), and then applied with Fsk, show a significant decrease in potentiation. Unpaired student’s t-test, p = 0.1236 (SC), p = 0.0055 (MPP) I_1,2_) PPR measurements show no change in PPR values following Fsk administration in the SC pathway. On the other hand, similar measurements in the MPP show slight decrease in PPR values to a similar degree across genotypes. Two-way repeated-measures ANOVA, p > 0.05 J_1,2_) PPR mean values 40 minutes after Fsk application for the two experimental groups. K_1,2_) Representative traces of paired-pulse recordings before (faint color) and 40 minutes after (bold color) Fsk administration for WT (floxed) (blue) and CaMKII-Rapgef2-cKO (red) conditions.

### CaMKII-Cre-Rapgef2-cKO mice have impaired Fsk-induced SC-LTP and Fsk-induced MPP-LTP

Since RapGEF2 is a cAMP sensor, we tested whether RapGEF2 is involved in cAMP-dependent synaptic potentiation in SC synapses. Accordingly, we compared hippocampal slices obtained from CaMKII-Rapgef2-cKO mice and their floxed littermates for their ability to undergo Fsk-induced SC-LTP. After establishing a stable baseline of fEPSPs, Fsk was perfused to the recording chamber for 5 minutes and then washed out using aCSF. This was followed by another 45 minutes of recording, during which the effect of Fsk was assessed. Our measurements of SC fEPSPs before and after Fsk application have yielded a small but significant increase in fEPSP amplitudes following Fsk application in the control group (floxed control, 118.6 ± 2.56). However, potentiation of cKO slices was insignificant 40 minutes after Fsk application (cKO, 103.7 ± 1.43; Figure 4E_1_-G_1_, two-way repeated-measures ANOVA, F_(1, 11)_ = 6.244, p = 0.0296, unpaired student’s t-test, p = 0.0002). These results imply that RapGEF2 is necessary for Fsk-induced potentiation in the SC-CA1 synapse. To elucidate the relevance of the MEK→Erk pathway to the observed RapGEF2-mediated Fsk induced potentiation, we tested whether U0126 application on control slices would phenomimic the effect of RapGEF2-KO. Consistent with a shared molecular pathway of RapGEF2, MEK and Erk, our results show that U0126 application abrogated Fsk-induced potentiation in control slices (floxed control, 147 ± 6.8; cKO, 112.4 ± 2.38; Figure 4H_1_, unpaired student’s t-test, p = 0.1236). This form of LTP was not accompanied by any change in PPR, suggesting that Fsk-LTP is expressed mainly in the postsynaptic compartment of the SC-CA1 synapse and is dependent on RapGEF2 (Figure 4I_1_-K_1_, two-way repeated-measures ANOVA, F_(1, 10)_ = 0.1727, p = 0.6865).

To determine whether RapGEF2 involvement in Fsk-induced synaptic potentiation is restricted to the SC-CA1 pathway, or takes place in other hippocampal synapses, we tested whether RapGEF2 is also involved in Fsk-induced LTP in entorhinal cortex to granule cell synapses (EC-DG synapse) through the medial perforant path (MPP). This synapse also exhibits postsynaptic LTP, is involved in various cognitive behavioral tasks, and shows reduction of RapGEF2 expression in CamK-Cre::RapGEF2^fl/fl^ mice. We measured the degree by which acute slices prepared from brains of either floxed control or cKO mice undergo MPP-LTP upon exposure to Fsk. For this purpose, acute hippocampal slices obtained from either floxed control (RapGEF2^fl/fl^) or cKO (CamK-Cre::RapGEF2^fl/fl^) littermates were used for extracellular field recordings from the MPP. While slices from floxed controls showed 52.7 ± 4.9% increase in fEPSPs amplitudes compared to their baseline levels, which lasted for over 45 minutes, slices prepared from CaMKII-RapGEF2-cKO mouse brains showed an initial increase in synaptic efficacy, which gradually decreased to significantly lower levels relative to floxed control mice (111.9 ± 4.4) after around 40 minutes (Figure 4E_2_-G_2_, two-way repeated-measures ANOVA, F_(1, 14)_ = 26.65, p = 0.0001, unpaired student’s t-test, p < 0.0001). In addition, as RapGEF2 might mediate Fsk-induced potentiation via the MEK/ERK pathway, we exposed slices derived from floxed control mice to U0126, and examined whether or not Fsk-induced potentiation was blocked. In agreement with the finding obtained for the SC-CA1 synapse, floxed control mouse hippocampal slices exposed to U0126 during the entire recording showed an abrogation of Fsk-induced potentiation at the MPP synapse, with reduction to levels seen in cKO mouse slices (Figure 4H_2_, unpaired student’s t-test, p = 0.0055). Hence this Fsk-induced potentiation is dependent on RapGEF2 and MEK kinase activity, suggesting that it is mediated by the MEK→ERK pathway. In addition, we monitored changes in PPR by delivering a paired-pulse stimulation (ISI of 40 milliseconds) via the stimulating electrode during these recordings. Application of Fsk was associated with a 10% decrease in the PPR in both floxed control and cKO mouse slices. These changes are consistent with a small increase in presynaptic release probability that was elicited by Fsk. However, these presynaptic modifications seem to be RapGEF2-independent, as cKO slices showed similar PPR levels at the MPP synapse following Fsk application (Figure 4I_2_-K_2_, two-way repeated measures ANOVA, F_(1, 14)_ = 4.154, p = 0.0609). Nevertheless, Fsk potentiation depends on RapGEF2/ERK as RapGEF2 depletion or U0126 application led to reduced Fsk-dependent potentiation. As the presynaptic effect was RapGEF2-independent, we suggest that RapGEF2 contributes to the postsynaptic Fsk-induced potentiation, which was reduced in RapGEF2 KO (cKO) slices. These results are consistent with the demonstration of a role for ERK in Fsk-induced postsynaptic LTP at the MPP synapse.

### Differential dependency on RapGEF2 in fear conditioning-induced immediate early gene activation

The dynamic changes of expression of immediate-early genes (IEGs), such as c-Fos and Egr-1, have been widely used as molecular markers for neuronal plastic changes measured in cell culture (Harada et al., 2001; Han et al., 2007; Adams et al., 2017) and in vivo ((Jiang et al., 2021) and references therein), especially in the context of genetic manipulation of hippocampal function required for memory formation (Liu et al., 2012; Ramirez et al., 2013; Liu et al., 2014; Gallo et al., 2018). Previous studies suggested that most neuronal ensembles encoding fear memory may simultaneously express various IEGs, however, to what extent IEG-positive ensembles overlap with each other and whether they are regulated separately in the same neurons, remains unclear. To examine the fear conditioning-induced expression changes of IEGs, particularly c-Fos and Egr-1, brain sections were obtained from floxed control and cKO mice 1 h after fear conditioning and were prepared for immunostaining (Figure 5). Quantification of resultant images indicated that c-Fos was significantly induced after fear conditioning (Figure 5A & 5B, two-way ANOVA followed by *post hoc* Bonferroni t-test to compare two genotypes with or without fear conditioning) throughout CA1 (F_(1,16)_=5.598, p=0.031), DG (F_(1,16)_=27.764, p<0.001) and CA3 (F_(1,16)_=12.774, p=0.003) in the hippocampus, and BLA (F_(1,15)_=25.832, p<0.001). However, RapGEF2 ablation in these areas had no effect on c-Fos expression (genotype effect: F_(1,16)_=0.0611, p=0.808 for CA1; F_(1, 16)_=0.0293, p=0.866 for DG; F_(1,16)_=0.106, p=0.749 for CA3; F_(1,15)_=0.00121, p=0.973 for BLA) (Figure 5A & 5B). On the other hand, the increase of Egr-1 expression induced by fear conditioning is RapGEF2-dependent in hippocampal CA1 and DG (Figure 5C & 5D). Similarly to c-Fos expression, Egr-1 expression also showed a fear conditioning effect in hippocampal CA1 (F_(1,16)_=21.01, p<0.001), DG (F_(1,16)_=14.788, p=0.001), CA3 (F_(1,16)_=5.03, p=0.039) and BLA (F_(1,16)_=21.577, p<0.001) (Figure 5D). However, with ablation of RapGEF2 in CA1 and DG, Egr-1 induction after fear conditioning was significantly attenuated in cKO mice only in CA1 and DG (genotype effect: F_(1,16)_=6.566, p=0.021 for CA1; F_(1, 16)_=6.161, p=0.025 for DG, with *post hoc* all pairwise multiple comparison in CA1: FC vs NFC within flox, p<0.001; flox vs cKO within FC, p=0.007, and in DG: FC vs NFC within flox, p<0.001; flox vs cKO within FC, p<0.001). Consistent with the maintenance of RapGEF2 expression in CA3 in the Camk2a-Cre^+/-^::RapGEF2^fl/fl^ mice, there were no genotype-specific IEG effects observed in CA3 (F_(1,16)_=1.229, p=0.284). No genotype effect was observed for Egr-1 expression in BLA (F_(1,16)_=0.163, p=0.692), confirming the involvement of the hippocampus, and not the BLA, in mediating the RapGEF2-dependent effects on contextual fear conditioning, but not cue fear conditioning, documented here. In addition, RapGEF2-dependent Egr-1 induction, but not c-Fos induction, after fear conditioning suggests that the pacellated signaling pathways involved in IEG induction. As Egr-1 is activated by ERK, and is absence in the hippocampus from RapGEF2 cKO mice, and as Fsk induced potentiation depends on RapGEF2 and ERK signaling, it strengthen the link between the involvement of RapGEF2 in Fsk-induced potentiation and memory fear conditioning. Further characterization of fear-conditiong-induced IEGs mRNA by RNAscope with egr-1 and c-fos specific probes indicated that upregulation of c-fos mRNA in hippocampal CA1 and DG reached a peak 30 minutes after fear conditioning (Figure 5E). Up-regulation of c-fos occurred mainly in neurons that also exhibited increased egr-1 mRNA, and most egr-1-positive neurons were also fos-positive. Thus, at this time point, 67.77% ± 10.92% of the egr-1 positive neurons were also c-fos positive in CA1, and 87.89% ± 7.67% of egr-1 positive neurons were also c-fos positive in DG (Figure 5E), indicating that the up-regulation of c-fos and egr-1 largely occurs in the same neuronal population(s) during fear conditioning, rather than in separate neuronal populations. This observation strongly supports our working hypothesis that while up-regulation of multiple immediate-early genes occurs in neurons upon activation during physiological events such as stress responding and fear conditioning, this up-regulation is parcellated through separate signaling pathways, in this case via a RapGEF2-dependent one for Egr-1, and a RapGEF2-independent one for c-Fos.

**Figure 5.**
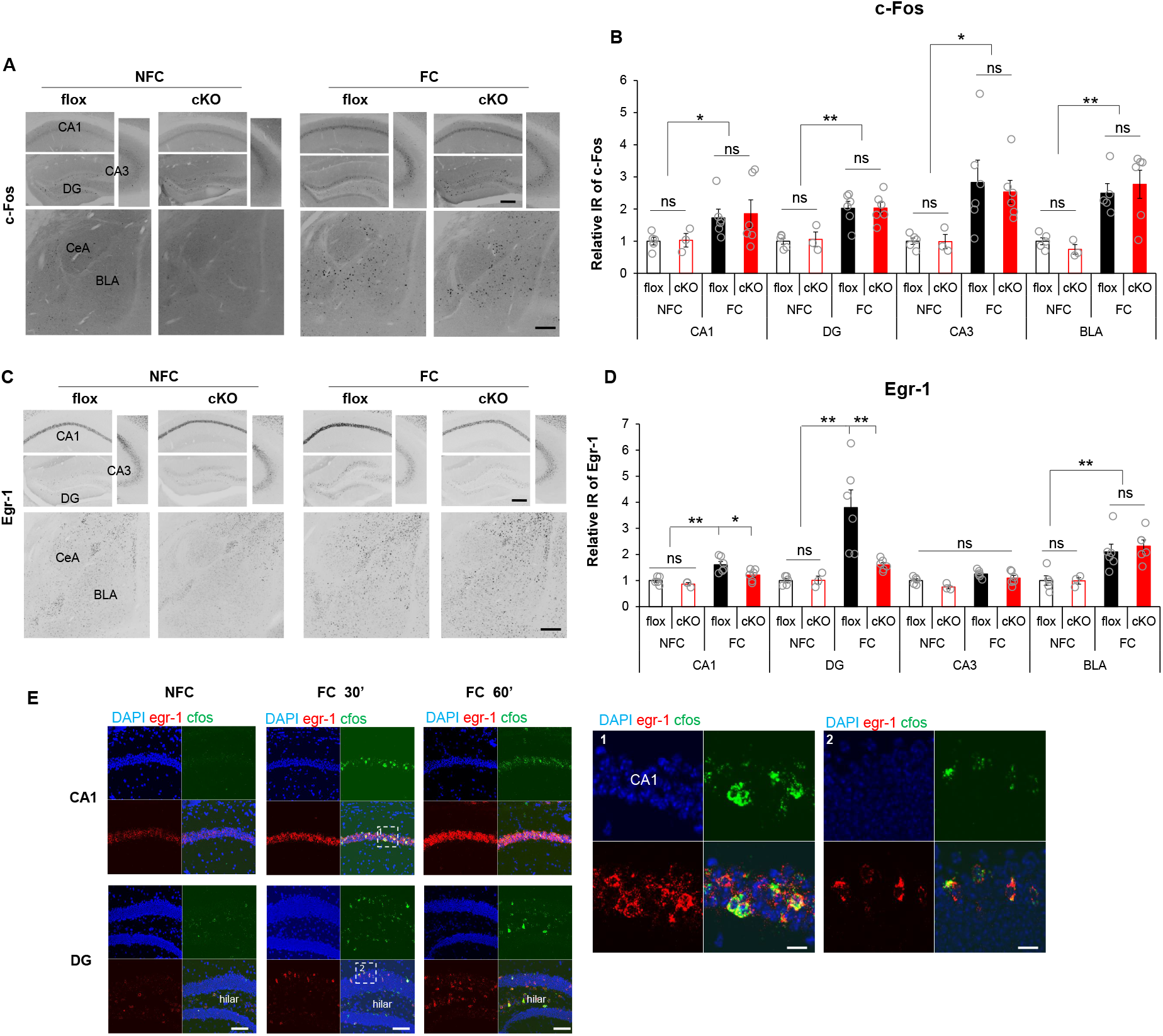
Differential dependency of RapGEF2 in fear conditioning-induced immediate early genes activation. (A) Representative images of cFos immunostaining in hippocampal CA1, DG and CA3 and basolateral amygdala of flox and cKO mice that were sacrificed 1hr after fear conditioning (FC) or stayed in the home cage (NFC). Scale bar: 200 μm (B) Quantification of cFos IR in hippocampal subregions and basolateral amygdala of flox and cKO mice indicated that fear-conditioning induced cFos increase in all these regions. No significant difference was observed between flox and cKO mice. N=3~6 for each group. Two-way ANOVA following by *post hoc* Bonferroni t-test, *p<0.05, **p<0.001. (C) Representative images of Egr-1 immunostaining in hippocampal CA1, DG and CA3 and basolateral amygdala of flox and cKO mice after fear conditioning. Scale bar: 200 μm (D) Quantification of Egr-1 IR indicated that fear-conditioning induced Egr-1 increase in CA1 and DG is RapGEF2-dependent. cKO mice with RapGEF2 ablation in CA1 and DG showed attenuated Egr-1 increase in CA1 and DG 1hr after fear conditioning, compared to flox mice. N=3~6 for each group. Two-way ANOVA following by *post hoc* Bonferroni t-test, *p<0.05, **p<0.001. (E) RNAscope with egr-1 (in red) and c-fos (in green) probes indicated that upregulation of c-fos mRNA in hippocampal CA1 and DG following fear conditioning occurred exclusively in the neurons with upregulation of egr-1 mRNA. In CA1, 67.77% ± 10.92% of Egr-1 positive neurons are c-fos positive. In DG, 87.89% ± 7.67% of egr-1 positive neurons are cfos positive. Scale bar: 100 μm (left panels), 20 μm (right panels).

## DISCUSSION

The data presented in this report provide evidence for the assertion that the cAMP→RapGEF2→ERK pathway is required for cAMP-dependent LTP and long-term memory (LTM) supporting contextual fear conditioning. This assertion is based on the dependence of contextual conditioned fear learning on RapGEF2; the dependence of learning-associated ERK activation on RapGEF2; the dependence of cAMP dependent synaptic potentiation on RapGEF2; and an extensive literature supporting a requirement for ERK in contextual conditioned fear learning mediated by the hippocampal tri-synaptic pathway (Atkins et al., 1998; Selcher et al., 1999; Adams and Sweatt, 2002; Athos et al., 2002; Sweatt, 2004; Shalin et al., 2006; Sindreu et al., 2007).

Synaptic potentiation has long been invoked as a potential mechanism underlying associative learning. In a variety of learning paradigms, and in a number of vertebrate and non-vertebrate species including mammals, behavioral adaptations based on experience are accompanied by altered neurotransmission at specific synapses or in specific brain areas. These alterations are thought to increase the likelihood that a cue associated with an unconditioned stimulus can elicit the response originally elicited by the unconditioned stimulus, in its absence (Lynch, 2004; Bromer et al., 2018). The cellular and molecular mechanisms of synaptic potentiation, in which a biochemical or electrical signal conveyed to a recipient neuron increases its response to a standard excitatory input, have been studied extensively both in vivo and ex vivo (Lynch, 2004).

Synaptic potentiation can be of short duration (STP) or long duration (LTP), and LTP itself possesses at least two distinct features. One aspect of LTP requires elevation of cAMP (Frey et al., 1993; Silva et al., 1998; Winder et al., 1999; Kandel, 2001; Morozov et al., 2003), and a second (Collingridge and Bliss, 1995; Bliss et al., 2018), without an explicit cAMP requirement, can be elicited by tetanic electrical stimulation of neuronal inputs, and mimicked by glutamate in ex vivo models of hippocampal LTP. Here, we show, using forskolin to initiate cAMP-dependent LTP, that the biochemical hallmarks of this form of long-term potentiation are consistent with its role in the maintenance of fear conditioning in mice. Since ERK dependency distinguishes cAMP-dependent from cAMP-independent synaptic potentiation (Morozov et al., 2003), we sought a molecular actor involved in cAMP-dependent activation of ERK, and then asked if deletion of this signaling molecule from hippocampal neurons might abrogate both cAMP-dependent ERK activation, downstream ERK-dependent immediate early-gene induction, and potentially cognate hippocampal-dependent memory functions such as associative learning.

Our rationale for consideration of RapGEF2 as such a candidate derives from our previous results that RapGEF2 is an obligate cAMP sensor in Gs-coupled GPCR initiation of neuritogenesis in neuroendocrine cells (Emery et al., 2013); that RapGEF2 is a specifically neuronal/neurendocrine signaling component in adult mammals (Jiang et al., 2017); that RapGEF2 is required for dopaminergic cAMP-dependent signaling to ERK underlying cocaine’s psychomotor stimulant effects on behavior (Jiang et al., 2021); and that RapGEF2-dependent signaling to ERK is required for cAMP-dependent induction of the immediate-early gene Egr1/zif268 (Jiang et al., 2017; Jiang et al., 2021), implicated in learning and memory mechanisms, along with ERK and cAMP, in hippocampus as well as other brain areas (Bozon et al., 2003).

The requirement for RapGEF2 for both cAMP-dependent LTP, and for context-dependent fear conditioning in mice, provides a biochemical mechanistic linkage between cAMP, activation of Rap1 which is also implicated in cAMP-dependent LTP at the Schaffer collateral (CA3-CA1) synapse (Morozov et al., 2003; Zhang et al., 2018), and ERK, and prompted us to further examine in detail immediate-early gene induction associated with fear conditioning in vivo. LTP involves the mobilization of several immediate-early genes activated by both the cAMP and calcium pathways. Fos up-regulation can be either calcium- or cAMP-dependent, and that of other immediate early genes, such as Egr-1/Zif268, are cAMP-dependent (Sheng and Greenberg, 1990; Sheng et al., 1990; Ginty et al., 1992; Greenberg et al., 1992; Tai et al., 2001; Bozon et al., 2002; Bozon et al., 2003; Ravni et al., 2008; Gangarossa et al., 2011; Jiang et al., 2021). More controversial is the mode of regulation of fos initiated either by cAMP or calcium: this has been attributed both to a PKA-dependent, Rap1-independent activation of ERK, leading to MSK-1 activation and phosphorylation/activation of the transcription factor CREB, or to a CREB-dependent mechanism due CREB phosphorylation directly by PKA (Wong et al., 1999; Poser and Storm, 2001; Sindreu et al., 2007; Xia and Storm, 2012). Our finding that c-Fos up-regulation after fear conditioning occurs within the same hippocampal neuronal population(s) in which Egr-1/Zif268 is also elevated, allowed us to test whether separate (parcellated) signaling pathways exist for the regulation of these two IEGs, or if both are regulated by a common signaling pathway. We show here that activation of ERK and Egr-1/zif268 in CA1 and dentate gyrus, consequent to fear conditioning are RapGEF2-dependent, while fos up-regulation is not, thus supporting the ‘parcellation of signaling’ hypothesis.

Establishment of linkage between hippocampal cAMP-dependent synaptic potentiation, ERK activation and associative learning via a common requirement for RapGEF2 opens several avenues for future research. First, it must be ascertained whether the RapGEF2 dependence for patent functioning of the trisynaptic pathway resides primarily within the Schaffer collateral or the perforant pathway synapse, or both. The present data set provides equal evidence for all three possibilities. However, persistent (24 hour) context-shock association is impaired in the absence of RapGEF2, and early (3 hour) context-shock association, is not. Impairment of early effects on context-dependent association have been taken by some to indicate discrimination deficits, more commonly associated with defects in dentate gyrus compared to CA1. For this reason, deficits at the latter synapse are more likely to be reflected in the context-dependent fear learning deficits reported here in hippocampally RapGEF2-deficient mice.

Second, we have not established whether Rap1 or Rap2 is required for the actions of RapGEF2 for ERK activation in cAMP-dependent LTP, immediate early gene activation after fear conditioning, or contextual fear conditioning itself. We have previously demonstrated, in the neuroendocrine NS-1 cell line, that RapGEF2 is linked to activation of Rap1, whereas the cAMP sensor Epac preferentially activates Rap2 (Emery et al., 2017). RapGEF2 has been previously invoked in hippocampal signaling: RapGEF2 is a required component in Plk2-dependent synaptic plasticity in the hippocampus (Lee et al., 2011), has been invoked in mir19-dependent regulation of the migration of nascent granule cells in dentate gyrus (Han et al., 2016), and is involved in hippocampal synaptic maintenance and cognitive function in a mouse model for Alzheimer Disease (Jang et al., 2021). However, the role of Rap isoforms in RapGEF2-associated synaptic events in the hippocampus remain unclear. Exploration of Rap isoform-dependence of RapGEF2 involvement in cAMP-dependent LTP, and contextual fear conditioning may shed further light on this point, as well as on RapGEF2 involvement in long-term depression, also implicated in hippocampusdependent fear conditioning (Ryu et al., 2008; Stornetta and Zhu, 2011; Zhang et al., 2018).

This work demonstrates that cAMP signaling to ERK in hippocampal neurons is mediated through RapGEF2. RapGEF2 is necessary for both cAMP-dependent synaptic potentiation, phosphorylation of ERK, induction of the immediate early gene egr-1/zif268 and for contextual fear conditioning. Hence, our findings suggest that the dependence on this novel plasticity signaling molecule will provide an important criterion for linkage of cAMP-dependent synaptic potentiation to specific learning-dependent behaviors in future work.

## ACKNOWLEDGEMENTS

This work was supported by NIMH-IRP Project MH002386 to L.E.E., supplemental NIMH-IRP funding to the Dendritic Dynamics Hub (L.E.E., and H.A.T.), grants from Israel Science Foundation (ISF grants 953/16 and 2141/20) and the DFG (NA: 207/10-1) and and the Taube/Koret Global Collaboration in Neurodegenerative Diseases to UA., and training grant, a Brain and Behavior Research Foundation NARSAD Young Investigator Award and supplemental DDH funding to M.S. Thanks to Haiying Zhang for mouse colony management and genotyping, and to the NINDS Light Imaging Facility for microscopy support.

